# The roles of Conserved Domains in DEMETER-Mediated Active DNA Demethylation *in planta*

**DOI:** 10.1101/175091

**Authors:** Changqing Zhang, Yu-Hung Hung, Xiang-Qian Zhang, Dapeng Zhang, Wenyan Xiao, Lakshminarayan M. Iyer, L. Aravind, Jin Hoe Huh, Tzung-Fu Hsieh

## Abstract

DNA methylation plays critical roles in maintaining genome stability, genomic imprinting, transposon silencing, and development. In Arabidopsis genomic imprinting is established in the central cell by DEMETER (DME)-mediated active DNA demethylation, and is essential for seed viability. DME is a large polypeptide with multiple poorly characterized conserved domains. Here we show that the C-terminal enzymatic core of DME is sufficient to complement *dme* associated developmental defects. When targeted by a native DME promoter, nuclear-localized DME C-terminal region rescues *dme* seed abortion and pollen germination defects, and ameliorates CG hypermethylation phenotype in *dme-2* endosperm. Furthermore, targeted expression of the DME N-terminal region in wild-type central cell induces *dme-*like seed abortion phenotype. Our results support a bipartite organization for DME protein, and suggest that the N-terminal region might have regulatory function such as assisting in DNA binding and enhancing the processivity of active DNA demethylation in heterochromatin targets.

---

Double fertilization during sexual reproduction in flowering-plants is a unique process that underlies the distinctive epigenetic reprogramming of plant gene imprinting. In the ovule, a haploid megaspore undergoes three rounds of mitoses to produce a 7-celled, 8 nuclei embryo sac that consists of egg, central, and accessory cells ^1^. During fertilization pollen grain elongates and delivers two sperm nuclei to the female gametophyte to fertilize the egg cell and the central cell, respectively. The fertilized egg cell forms the embryo that marks the beginning of the subsequent generation. Fertilization of the central cell initiates the development of endosperm that accumulates starch, lipids, and storage proteins and serves as a nutrient reservoir for the developing embryo ^2^, ^3^. Endosperm is the major tissue where gene imprinting takes place in plant. Genomic imprinting is the differential expression of the two parental alleles of a gene depending on their parent-of-origin, and is an example of inheritance of differential epigenetic states. In Arabidopsis, MET1-mediated DNA methylation and DME demethylation are two modes of epigenetic regulation critical for imprinted expression of many genes ^4, 5, 6, 7, 8^. For example, DEMETER (DME) is required for the expression of *MEA*, *FIS2*, and *FWA* in the central cell and in the endosperm while MET1 is responsible for the silencing of *FIS2* and *FWA* paternal alleles ^4^,7. Gene imprinting is essential for reproduction in Arabidopsis, and seeds that inherit a maternal *dme* allele abort due to failure to activate *MEA* and *FIS2*, essential components of the endosperm PRC2 complex required for seed viability, in the central cell ^4, 9^.

*DME* encodes a bifunctional 5mC DNA glycosylase/lyase required for active DNA demethylation in the central cell and the establishment of endosperm gene imprinting in Arabidopsis ^5^. Additionally, paralogs of DME, REPRESSOR OF SILENCING 1 (ROS1), DML2, and DML3 are required to counteract the spread of DNA methylation mediated by the RNA-directed DNA methylation (RdDM) machinery into nearby coding genes ^10, 11^. The three regions in the C-terminal half of DME protein (the A, Glycosylase, and the B regions, or as the AGB region hereafter) are conserved among the DME/ROS1 DNA glycosylase clade, and are required for DME 5mC excision activity *in vitro*. Thus, the AGB region comprise the minimal catalytic core for the enzymatic function, catalyzing direct excision of 5mC from DNA and initiating active DNA demethylation that influences transcription of nearby genes ^5, 9, 12^.

In Arabidopsis, DME-mediated DNA demethylation is preferentially targeted to small, AT-rich, and nucleosome-poor euchromatic transposons that flank coding genes ^13^. Consequently, demethylation in the central cell influences expression of adjacent genes only in the maternal genome, and is a primary mechanism of gene imprinting in plant ^5, 13, 14, 15^. In addition to small TEs near coding sequences, DME also targets gene-poor heterochromatin regions for demethylation ^13^. The mechanism of DME recruitment to its target sites is not known. Studies in ROS1 have uncovered several players required in the ROS1 demethylation pathway ^16, 17, 18^. Among them *IDM1* encodes a novel histone acetylase that preferentially acetylates H3K18 and H3K23 *in vitro*, and ROS1 target loci are enriched for H3K18 and K23 acetylation *in vivo* in an IDM1-dependent manner ^19^. Thus, IDM1 marks ROS1 target sites by acetylating histone H3 to create a permissible chromatin environment for ROS1 function. The Arabidopsis SSRP1 (STRUCTURE SPECIFIC RECOGNITION PROTEIN1), a component of the FACT (facilitates chromatin transcription/transaction) histone chaperone complex, has been shown to regulate DNA demethylation and gene imprinting in Arabidopsis ^20^. Linker histone H1 functions in chromatin folding and gene regulation ^21, 22, 23, 24^, and was shown to interact with DME in a yeast two-hybrid screen and in an *in vitro* pull-down assay ^25^. Loss-of-function mutations in *H1* genes affect the imprinted expression of *MEA* and *FWA* in Arabidopsis endosperm, and impair demethylation of their maternal alleles, suggesting that H1 might participate in the DME demethylation process by interaction with DME ^25^.

Computational analysis showed that the DME/ROS1 like DNA glycosylases contain a core with multiple conserved globular domains, and except for the well-characterized glycosylase domain, very little is known about the function of the other domains. Here we show that the C-terminal region of DME necessary for 5-methylcytosine excision activity *in vitro* is sufficient to complement *dme* seed abortion and pollen germination defect, and partially rescue DNA hypermethylation phenotype in endosperm. We present evidence that the region N-terminal to the glycosylase domain can affect endogenous DME activity in a dominant negative manner when ectopically expressed in the nuclei of wild-type central cells. We propose a bipartite structural and functional organization model for the DME/ROS1 family of DNA glycosylases consisting the modular C-terminal AGB region that can substitute for DME’s developmental function and the NTD region that might have regulatory functions such as assisting DNA binding and enhancing the processivity of demethylation in heavily methylated genomic regions.

## Results

### The DME catalytic core region is sufficient to complement *dme* associated developmental defects

Previous studies have revealed that the C-terminal half of DME comprising the three conserved <u>A</u>, <u>G</u>lycosylase, and <u>B</u> regions (the <u>AGB</u> region, as shown in Supplementary Fig. 1a) are required for *in vitro* 5mC excision activity ^5^, and deletion of the non-conserved linker between domain A and the glycosylase domain (interdomain 1; ID1) does not affect DME *in vitro* enzymatic activity ^26, 27^. Thus, the AGB region is thought to be the core catalytic region for DME *in vitro* enzymatic activity. However, it is unknown whether the AGB region alone is sufficient for DME function *in vivo*. To determine if the AGB region is functional *in vivo*, we tested if expressing the AGB region in the central cell can complement *dme* seed abortion phenotype. A transgene carrying a 3.1-kb *DME* cDNA that encodes the C-terminal half of DME (DME^CTD^, residue 936-1987) under the control of a native DME promoter was introduced into *DME/dme-2* heterozygous plants by using the floral dipping method ^28^. Since DME^CTD^ lacks a nuclear localization signal (data not shown), a classical SV40 nuclear localization signal (PKKKPKV) was introduced in front of the C-terminal fragment (designated as *nDME*^*CTD*^, see Supplementary Fig. 1b) to ensure proper nuclear localization. We obtained multiple independent transgenic lines and assessed the transgene’s ability to complement *dme-2* seed abortion phenotype.

The self-pollinated *DME/dme-2* plants produce 50% of normal seed that inherited wild type DME maternal allele, and the other 50% of aborted seed that inherited mutant *dme-2* maternal allele. In self-pollinated transgenic plants that carry a single locus of *nDME*^*CTD*^ or *DME*^*FL*^ (full length DME.2 cDNA, major isoform of DME ^29^) transgenes, we observed about 25% aborted seeds among independent transgenic lines, indicating that *nDME*^*CTD*^ and *DME*^*FL*^ complement *dme* seed abortion phenotype (Fig. 1a, b, Supplementary Table 1). In addition, we also transformed *nDME*^*CTD*^ and *DME*^*FL*^ into *dme-2/dme-2* homozygous plants (see Materials and Methods for isolation and characterization of *dme-2/dme-2* homozygous lines in Col-*gl*), both constructs produced T1 transgenic plants that displayed expected 50% seed abortion rate (Fig. 1b, Supplementary Table 1). Seed abortion caused by *dme* mutations is in part due to defects in activating imprinted PRC2 subunit genes required for endosperm development ^5, 9, 30, 31, 32^. We use qRT-PCR to check if nDME^CTD^ also restores DME target genes expression in the central cell. Indeed, *FIS2* and *FWA* expression is restored in the complemented lines (Fig. 1c). Thus nDME^CTD^ can substitute for the endogenous DME activity for seed viability, and active DME target genes expression.

**Figure 1.**
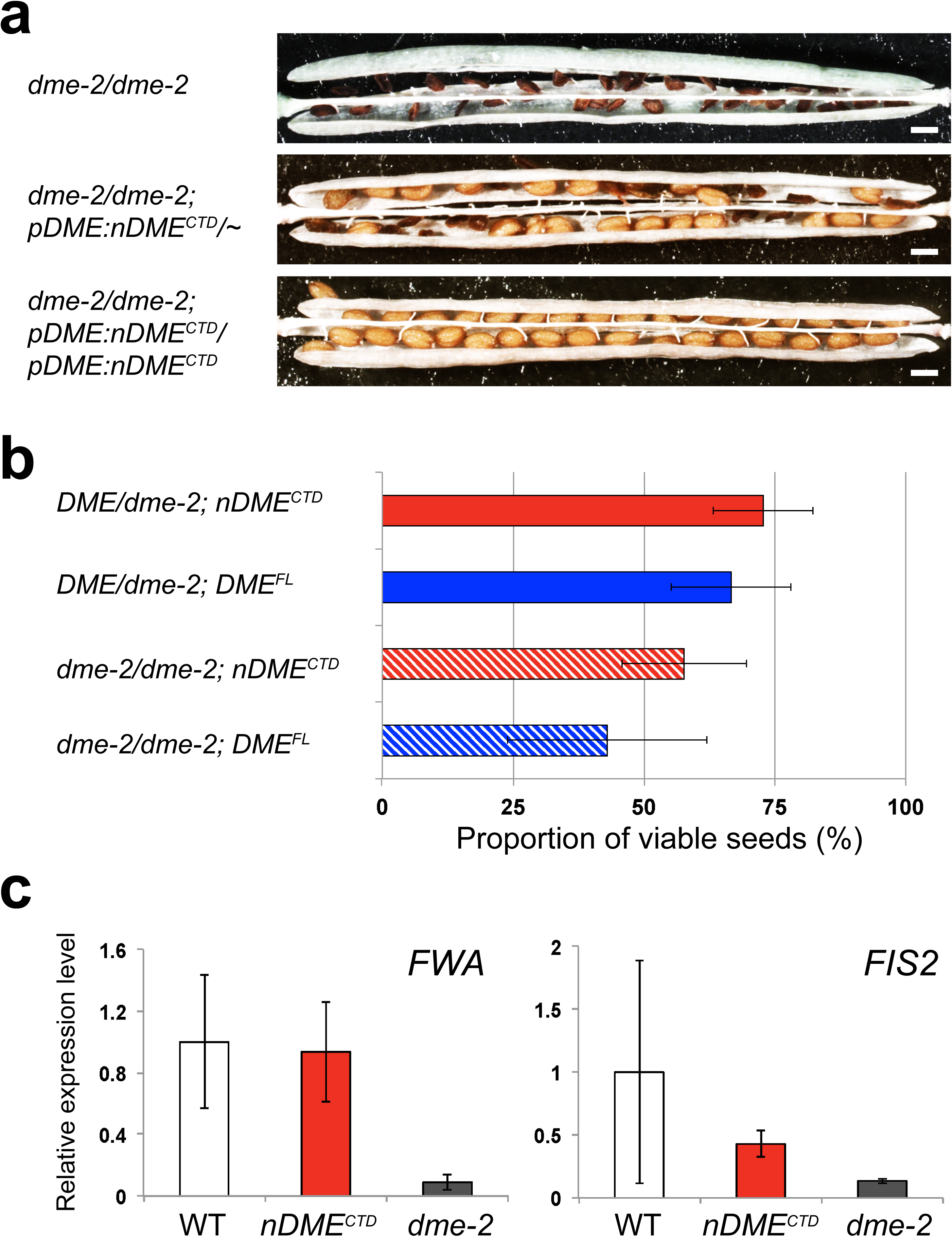
Complementation of *dme* seed abortion phenotype by the truncated DME nAGB. (a) Siliques were dissected and photographed 14 days after self-pollination. In *dme-2/dme-2* silique greater than 99% of seeds are aborted. A single copy of *nDME*^*CTD*^ transgene reduces seed abortion rate to 50%; and in the *dme-2/dme-2; nDME*^*CTD*^*/nDME*^*CTD*^ silique, all the *dme-2* seeds are rescued and developed normally. Scale bar = 0.5 mm. (b) Complementation of *dme-2* seed abortion phenotype by *nDME*^*CTD*^ *and DME*^*FL*^. (c) The *nDME*^*CTD*^ transgene restores DME target genes *FWA* and *FIS2* expression. WT: Col-0; *nDME*^*CTD*^: *dme-2/dme-2; nDME*^*CTD*^*/ nDME*^*CTD*^; *dme-2*: *dme-2/dme-2*. Total RNA was isolated from stage F1 to F12 floral buds.

In addition to maternal effects on seed viability ^9^, mutations in DME also affect pollen function in Col-0. When *DME/dme-2* heterozygous plants are self-pollinated, only about 20-30% of the viable F1 progeny are heterozygous (Supplementary Table 2), due to decreased *dme* pollen germination rate ^33^. To test whether nDME^CTD^ can rescue *dme* pollen phenotype, we pollinated wild type Col-0 with pollen derived from transgenic lines that are homozygous for the *dme-2* allele and carry a single locus of the *nDME*^*CTD*^ transgene (*dme-2/dme-2; nDME*^*CTD*^*/∼*). If nDME^CTD^ does not complement *dme-2* pollen germination defects, we expect roughly half of the F1 progeny will carry the nDME^CTD^ transgene (hygromycin resistant) because mutant pollen with or without the transgene would germinate with equal frequency. Instead, we observed 65% - 90% of the F1 progeny are hygromycin resistant (Table 1), indicating that nDME^CTD^ complements *dme-2* pollen germination defect. These results show that expressing the C-terminal half of DME protein in the nucleus is sufficient to rescue *dme* visible phenotypes *in planta*.

### nDME^CTD^ partially rescue *dme-2* CG hypermethylation phenotype in the endosperm

In Arabidopsis seed viability depends on the DME activity in the central cell to activate the MEDEA/FIS2/MSI1/FIE PRC2 complex required for endosperm development. In addition, DME is required to demethylate multiple maternally (*MEGs*) or paternally expressed imprinted genes (*PEGs*) to establish their parent-of-origin specific expression patterns in the endosperm ^13^,15. Thus, in *dme* mutant endosperm, discrete genomic loci targeted by DME for demethylation are hypermethylated ^13^. Since nDME^CTD^ complements *dme* seed abortion, and activates DME target gene expression (Fig. 1), we assumed it does so by demethylating the central cell genome and activating PRC2 genes essential for seed development. To test this hypothesis, and to examine the extend of nDME^CTD^ demethylation activity *in vivo*, we manually isolated *nDME*^*CTD*^*-* complemented endosperm (*dme-2/dme-2;nDME*^*CTD*^*/nDME*^*CTD*^), determined the DNA methylation profile by whole genome bisulfite sequencing, and compared the complemented methylomes to those of wild-type and *dme-2* endosperm. Methylomes from three independent lines were generated and compared with that of *dme-2* endosperm. We observed although the differentially methylated regions (DMRs) between each independent lines do not completely overlap, the DMRs unique to each line are also demethylated in other lines (Supplementary Fig. 2, 3), suggesting that the number of overlapped DMRs was underestimated due to the cutoff used in defining the DMRs, similar to what’s observed in a recent study ^34^. We therefore used the combined reads from three independent lines for the subsequent analyses so that all comparisons are confined to the same cutoff criteria (see Materials and Methods). As expected, several DME regulated *MEGs* and *PEGs* are demethylated compared to *dme-2* endosperm, indicating that nDME^CTD^ is correctly recruited to these loci for demethylation (Fig. 2a). We focused our analysis on previously determined differentially methylated sites between *dme-2* and wild-type endosperm (*dme* hyper-DMRs, the DME canonical targets) ^13, 15^. Overall, the CG methylation levels in these canonical DME target sites are reduced in the complemented endosperm, indicating that nDME^CTD^ is directed to these endogenous DME target sites for demethylation. However, compared to wt endosperm, these *dme* hyper-DMRs are demethylated to a lesser degree by the nDME^CTD^ (Fig. 2b). Thus nDME^CTD^ only partially rescues the *dme* CG hypermethylation phenotype in the endosperm. The DMRs of *dme* relative to wild-type endosperm or to *nDME*^*CTD*^-complemented endosperm partially overlap (Supplemental Fig. 4). However, among the DMRs unique to nDME^CTD^, we also observed decreased CG methylation in WT endosperm compared to *dme*, indicating that they are also demethylated by the endogenous DME. Similarly, among the DMRs unique to wt endosperm, these regions are also demethylated by the nDME^CTD^. Thus nDME^CTD^ appears to partially demethylate the majority of the loci targeted by the endogenous DME. These observations also suggest that intact full-length DME protein is required for robust and complete demethylation *in vivo*.

**Figure 2.**
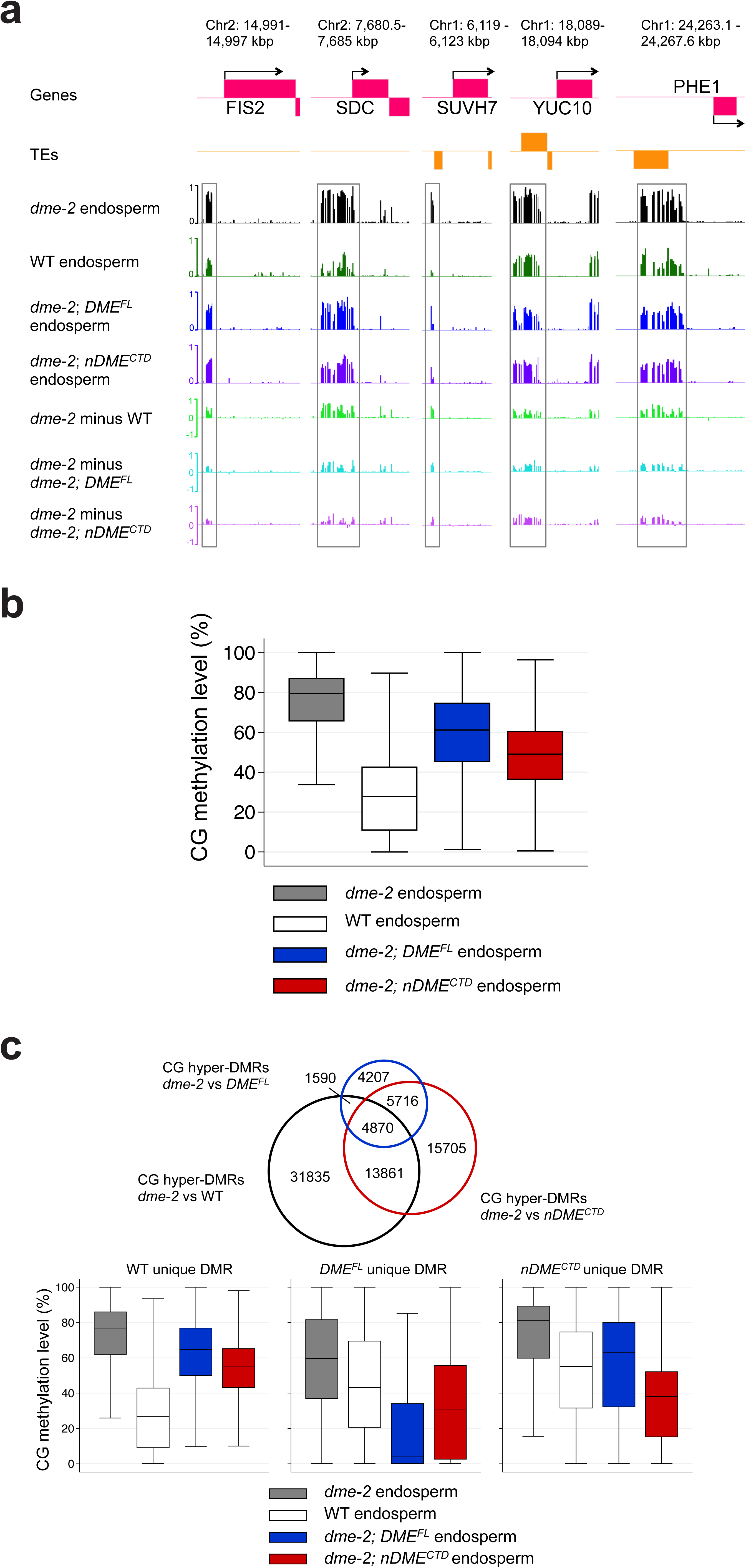
Endosperm methylome analysis. (a) Genome browser snapshots of CG DNA methylation at selected imprinted gene loci. Top two tracks are coding genes (magenta) and TEs (orange) with Tair10 chromosome coordinates. For the bottom seven tracks, each track represents fractional CG methylation levels for different genotype: black trace, *dme-2* endosperm; dark green trace, WT endosperm; dark blue trace, *DME*^*FL*^-complemented endosperm; dark purple trace, *nDME*^*CTD*^-complemented endosperm; light green trace, WT endosperm subtracted from *dme-2* mutant endosperm; light blue trace, *DME*^*FL*^-complemented endosperm subtracted from *dme-2* endosperm; light purple trace, *nDME*^*CTD*^-complemented endosperm subtracted form *dme-2* endosperm. DNA CG hypomethylation at selected maternally expressed (*FIS2* and *SDC*) and paternally expressed (*SUV7*, *YUC10, and PHE1*) imprinted genes is restored in *DME*^*FL*^-and *nDME*^*CTD*^-complemented endosperm. (b) Boxplot of CG methylation levels among canonical DME target sites in *dme-2* mutant (grey), WT (white), *DME*^*FL*^*-* (blue), or *nDME*^*CTD*^ *-* (red) complemented endosperm. (c) Venn Diagram (top panel) of CG hyper-DMRs in 50-bp windows between *dme-2* endosperm relative to WT, *DME*^*FL*^-complemented or *nDME*^*CTD*^-complemented endosperm. Boxplot (bottom panel) of CG methylation levels in *dme-2* mutant (grey), WT (white), *DME*^*FL*^*-* (blue) or *nDME*^*CTD*^*-* (red) complemented endosperm in WT only (left panel), *DME*^*FL*^ only, or *nDME*^*CTD*^ only (right panel) DMRs.

We next examined the methylome of *dme-2* endosperm complemented by the full length *DME.2 cDNA* (designated as *DME*^*FL*^). Unexpectedly, based on the number of DMRs between *dme* and *DME*^*FL*^-complemented endosperm and the level of CG methylation within the DMRs (Fig. 2c), DME^FL^ appears to be less active compared to endogenous DME, or to nDME^CTD^, albeit it being able to complement *dme* seed abortion (Fig. 1b) ^9, 35^. Since the *DME*^*FL*^ transgene only differs from *nDME*^*CTD*^ by the N-terminal region, reduced activity of DME^FL^ compared to DME^CTD^ cannot be attributed to the lack of introns or 3’ flanking sequences that might be needed for robust DME protein production. Indeed, we found both transgenes are expressed at comparable levels in *DME*^*FL*^*-* and *nDME*^*CTD*^-complemented lines used in the methylome study (Supplemental Fig. 5), indicating lower activity of DME^FL^ compared to nDME^CTD^ is not due to their differential transcript abundance. Nevertheless, comparison of CG methylation levels in DMR regions unique to DME^FL^, nDME^CTD^, or endogenous DME also reveals that unique DMR regions are more or less hypomethylated in WT or in complemented endosperm relative to *dme* endosperm. Thus the methylome difference between wt, *DME*^*FL*^*-*, and *nDME*^*CTD*^-complemented endosperm appears to be more in the degree of demethylation, rather than in targeting specificity.

### Function of the N-terminal region in DME-mediated active DNA demethylation

The *dme-2* allele is caused by an activation-tagging T-DNA insertion in the middle of the A region (Supplementary Fig. 1a)^9^. We found that in floral buds of *dme-2/dme-2* plants, the endogenous *DME* transcripts downstream of T-DNA insertion site is greatly reduced compared to wild-type Col-0 plants, but the level of DME transcripts upstream of the T-DNA insertion site is relatively high (Supplementary Fig. 6). We suspected these transcripts could produce truncated form of DME proteins that might interfere with the DME^FL^ transgene activity. To test this hypothesis, we transformed wild-type Col-0 plants with an engineered GFP-tagged DME NTD (using the genomic DNA sequence upstream of T-DNA insertion site, encoding residues 1-1022, designated as *DME*^*NTD*^*-GFP*) transgene mimicking the *dme-2* T-DNA insertion (Supplementary Fig. 1B). Clear GFP signals are observed in the central cell nuclei of transgenic lines (data not shown). We also observed about one third of transgenic lines showing apparent *dme-2* like seed abortion phenotype, with abortion rates ranging from 10% to ∼ 40% (Supplementary Table 3, 4) in the T1 plants, suggesting that expression of DME^NTD^ has a dominant negative effect on endogenous DME protein.

To minimize the possibility and the degree of transgene induced sense co-suppression, we reverse translated DME^NTD^ protein sequence into cDNA sequence using the human codon usage table. As a result, the re-engineered “humanized” version of NTD (mDME^NTD^) codes for the identical protein sequence but with no significant nucleotide sequence similarity to the original cDNA sequence to induce co-suppression (Supplementary Table 5). In addition, a GFP tag was added to the C-terminus (mDME^NTD^-GFP) to monitor its expression (Fig. 3a). We generated 28 independent transgenic lines, and among them 16 lines showed seed abortion rate of 5% - 52% (Supplementary Table 3, 6). The aborted seeds resemble *dme* mutant seeds with abnormal endosperm, arrested embryo, and shriveled brown seeds (Fig. 3b, c). We selected four lines with high, medium, or no seed abortion rate (Fig. 3d), and assessed the endogenous DME transcript abundance. As shown in Fig. 3e, among lines with different seed abortion rate, the endogenous DME mRNA abundance is similar to that of the vector control line, indicating the severity of seed abortion phenotype is not due to interference of endogenous *DME* transcripts. Furthermore, the rate of seed abortion is positively correlated with the levels of *mDME*^*NTD*^*-GFP* mRNA (Fig. 3f), suggesting the degree of seed abortion is likely due to the levels of transgene expression. We next tested whether expression of *nDME*^*CTD*^ or *DME*^*FL*^ in WT Col-0 can also induce seed abortion phenotype. For each construct, more than 25 independent transgenic lines were examined and none resulted in any seed abortion phenotype (Supplementary Table 3). Thus the dominant negative effect appears to be specific to the DME NTD region.

**Figure 3.**
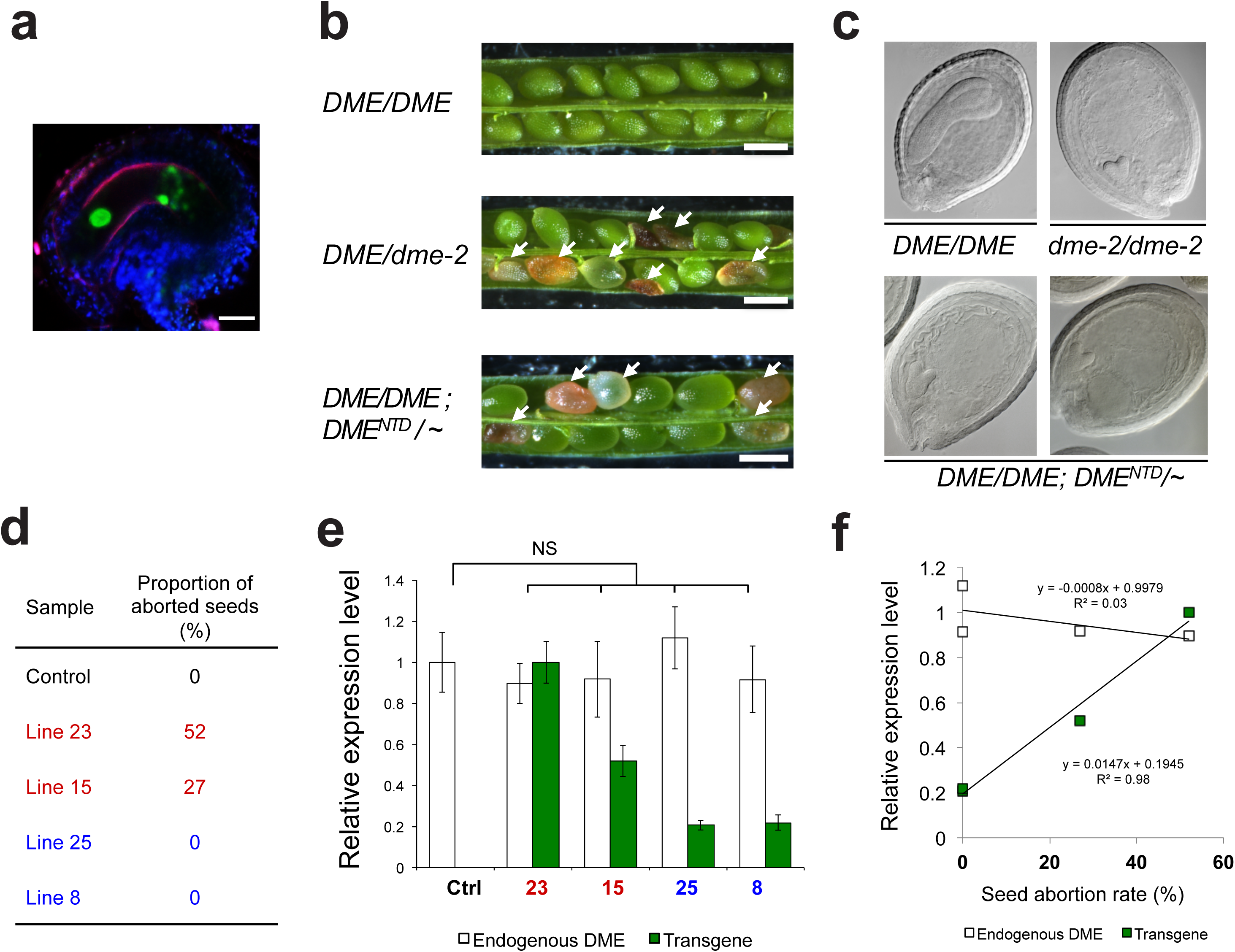
Expression of DME NTD region in wild-type central cell induces *dme*-like seed abortion phenotype. **(a)** Confocal microscopy image of ovule in F12 floral bud shows the expression of mDME^NTD^-GFP in the central cell. Scale bar, 20 μm. (**b-c**) Ectopic expression of *DME*^*NTD*^ in WT central cell induces *dme-2* like seed abortion phenotype in silique (**b**) and in developing seeds (**c**). Total RNA was isolated from stage F1 to F12 floral buds from independent lines with different seed abortion ratios (**d**) to assess transgene and endogenous DME expression. (**e**) Endogenous DME transcript levels in independent transgenic lines are comparable to the control line, but the transgene expression level varies among these independent lines with different seed abortion rates. Error bars indicate SD. NS, p >0.2 (Ctrl vs 23), p > 0.5 (Ctrl vs 15), p > 0.3 (Ctrl vs 25), p > 0.4 (Ctrl vs 8), not significant (two-tailed t test). (**f**) Correlation analysis shows that the transcript abundance of the transgene, but not that of the endogenous DME transcripts, correlates with seed abortion rates (by linear regression).

### Evolutionary history and late acquisition of the N-terminal region of DME-like proteins

We show the C-terminal half of DME is sufficient to complement *dme* mutant developmental phenotypes, and can be recruited to most of the DME target loci. Thus the DME^CTD^ most likely contains intrinsic targeting information. To gain insights from the evolution of the conserved domains in DME, we conducted sequence searches of the NR database with various homologs as query. The core of the DME-like proteins, as previously reported ^36^, comprises the catalytic glycosylase domain of the HhH (<u>h</u>elix-<u>h</u>airpin-<u>h</u>elix) modules followed by the FCL ([<u>F</u>e4S4] <u>c</u>luster <u>l</u>oop) motif and a divergent version of an RRM (<u>R</u>NA <u>R</u>ecognition <u>M</u>otif) fold domain (Fig. 4). The DNA glycoslase and FCL domains span the A and G regions, whereas the RRM fold domain corresponds to the B region of angiosperm DME homologs. A diversity of domains associate with the basic DME core can be found across various clades. Land plants and charophytes (Streptophyta) possess a permuted divergent version of the umethylated CpG recognizing CXXC domain (containing only one of two structural repeats of the classical CXXC domain) between the FCL and RRM domains. By contrast, one or more copies of the CXXC domain can be found in chlorophyte and stramenopile algae at distinct positions. Some algal DME homologs (from Chlorophyte and stramenopile) also possess other chromatin-modification reader (Tudor and PHD domains), DNA binding (AT-hook motif), and the DnaJ domain which interacts with the chaperone Hsp70 ^36, 37^. These accessory domains suggest a potential role for regulating the associated DNA glycosylase activity according to the DNA methylation (via CXXC) or chromatin status (via PHD, Tudor) of the cell in which they are expressed.

**Figure 4.**
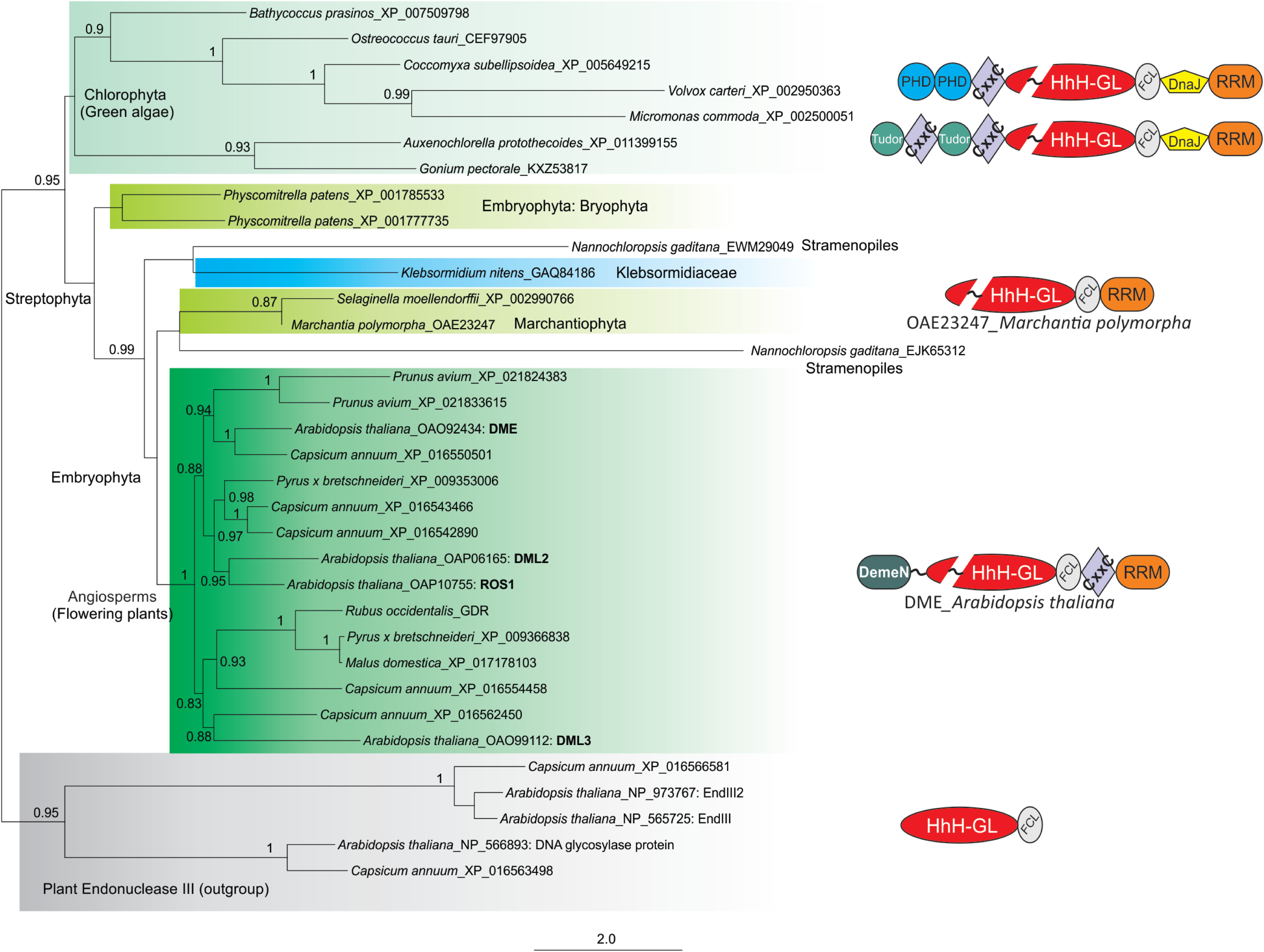
Evolution of plant DME-like proteins. A phylogenetic tree was reconstructed using the PhyML program. Only node supporting values >0.80 from ML bootstrap analyses are shown. The representative domain architectures of DME homologs in major plant clades are shown along the tree, demonstrating domain fusions during evolution. Domain abbreviations: DemeN, N-terminal domain of DEME-like proteins in angiosperms; DnaJ, DnaJ molecular chaperone homology domain (Pfam: PF00226); FCL, [Fe4S4] cluster loop motif (also called Iron-sulfur binding domain of endonuclease III; Pfam: PF10576); HhH-GL, HhH-GPD superfamily base excision DNA repair protein (Pfam: PF00730); PHD, PHD finger (Pfam: PF00628); RRM, RNA recognition motif (Pfam: PF00076); Tudor, Tudor domain (Pfam: PF00567).

The N-terminal half of the DME consists of a large portion of unstructured, low complexity sequences (residues 346-947), a stretch of basic amino acid-rich direct repeats (residues 291-345), and a 120 amino-acid N-terminal domain (DemeN) of unknown function (residues 1-120)(see Supplementary Fig. 7 for sequence alignment). The DemeN domain and charged repeats are restricted to the angiosperm lineage and appears to be a late acquisition during land plant evolution.

In summary, the evolutionary history of the DME domains can be summarized as follows: bacterial versions of the HhH-FCL pair from a cyanobacterial source fused to an RRM-fold domain and further acquired an insert in the glycosylase domain to give the ancestral form in the plant lineage. This was likely then transferred to the stramenopiles from a secondary chlorophyte endosymbiont of this lineage. Finally, at the base of the streptophyte radiation, DME acquired a permuted CXXC, and later the DemeN domain and the associated charged repeats were acquired in the angiosperm lineage, possibly to facilitate and ensure a robust and thorough demethylation.

## Discussion

We show for the first time that the core conserved region of the DME protein containing the DNA-glycosylase, FCL, divergent and permuted CXXC and divergent RRM domains is sufficient to rescue visible phenotypic defects caused by *dme* mutation. Although this truncated form of DME protein demethylates the majority of the canonical DME target sites, it does so in a less active and less efficient manner compared to the endogenous protein. We see two possibilities that might explain this lower activity and efficiency: 1) Critical cis-elements residing within introns or in 3’-end flanking sequences that are missing in the transgene might be required for robust transgene expression. 2) The N-terminal region might be required for full DME activity *in vivo*. Unfortunately, our attempt to assess the difference between DME^FL^ and nDME^CTD^ was confounded by the possible interference from truncated NTD proteins due to T-DNA insertion in *dme-2* background. We suspect this might contribute to the reduced DMRs observed in *DME*^*FL*^complemented endosperm. Therefore, we believe it is premature to draw any conclusion based on direct comparisons between *DME*^*FL*^-and *nDME*^*CTD*^-complemented endosperm methylomes (Fig. 2c).

Since the C-terminal AGB region is sufficient for DME’s seed viability function in *planta*, and can be recruited to most of the canonical DME target sites, the CTD polypeptide most likely contains sufficient targeting information. *in vitro* studies of ROS1 suggest that the B region containing the CXXC and RRM domains is essential for the glycosylase and lyase activities, and might recognize modified DNA^38^. It is possible that the permuted CXXC domain is required to direct the protein to the target sites, or is involved in discriminating methylated vs un-methylated cytosines^39^. This is supported by mutation studies that implicate a potential role for this domain in DME *in vivo* function, but not in vitro enzymatic activity (Huh and Hsieh, unpublished results). Similarly, the role of the enigmatic divergent RNA-recognition motif (RRM) domain is also not fully understood. Mutagenesis screens for residues required for demethylation activity in bacteria identified multiple amino acid residues within the RRM domain^40^. Although the involvement of RNA species in the active DNA demethylation process has not been firmly established, an RRM protein ROS3 required for ROS1 demethylation suggests a potential role of non-coding RNAs in the active DNA demethylation pathway in Arabidopsis^41^. While it is tempting to speculate a role for RNA-binding, the DME RRM might also bind single-stranded DNA with methylated bases.

Based on the reduced demethylation activity of nDME^CTD^ on the canonical DME target sites, we suspect the NTD region might be required for full and robust demethylation activity probably to ensure that the imprinting network is properly activated and maintained (e.g., by subsequent PRC2 activity). To achieve this, the DME NTD might function to assist the glycosylase enzyme by tightly binding to DNA template for more complete and thorough demethylation. Supporting such model, *in vitro* study of ROS1 activity on 5mC excision revealed that the basic repeats (3DR, AT-hooks) region binds strongly to DNA template non-specifically, and removal of NTD region impairs the sliding capacity of the protein on DNA template^42^, and significantly reduced ROS1 5mC excision activity^43^. We observed reduced degree of demethylation by nDME^CTD^ regardless of target length (Supplemental Fig. 8), suggesting that NTD is needed for complete demethylation in all the target sites.

Although DME preferentially targets smaller euchromatic transposons that flank coding genes, it also targets gene-poor heterochromatin regions for demethylation ^13^. The biological significance of heterochromatin demethylation by DME is not known, but was speculated to involve reinforcing DNA methylation in egg cell and subsequently in the embryo ^13^. These heterochromatin target sites are densely methylated, and demethylation by DME results in longer DMRs between *dme-2* and wt endosperm. Interestingly, the number of longer DMRs is significantly reduced between *dme-2* and *nDME*^*CTD*^-complemented endosperm, suggesting that removal of NTD region also reduces the processivity of demethylation in long target sites (Supplemental Fig. 9a). Since heterochromatin regions are compacted, demethylation in such loci will require substantial chromatin remodeling such as eviction of nucleosomes for DME to gain access to the templates. It is tempting to speculate that the conserved motif in the DemeN domain might recruit other factor(s) via protein interaction to remodel local chromatins to permit DME demethylation. However, based on current data we cannot unequivocally ascribe NTD’s function due to lack of proper full length DME transgenic comparison. Nevertheless, our results caution that peculiarity in certain genetic backgrounds (e.g., *dme-2*) might confound data interpretation. Future work on DME functional study could benefit from the generation of a clean loss-of-function background such as deleting the entire DME locus using CRISPR-assisted genome editing techniques.

We envision a possible model where the AGB region is sufficient for directing DME to target loci while NTD region is required for interacting with local chromatin environment, stabilizing binding to chromosomal templates, and assisting demethylating flanking sequences. In the absence of NTD, nDME^CTD^ can still demethylate majority of target sites, but in a less-efficient manner, likely due to the lack of non-specific DNA-binding by the basic AT-hook motifs. We surveyed wt DMRs that are longer than 1.5 kb, and found that these regions are also nDME^CTD^’s DMRs, but are shorter in length (Supplemental Fig. 9b), possibly due to missing the DemeN domain. If NTD is needed for longer and more robust demethylation, why ectopic expression of NTD causes dominant negative (DN) effects on endogenous protein? Classical examples of dominant negative mutation often involve protein-protein interactions that are disrupted by mutated or truncated form of one particular partner or subunit. Although we do not have any evidence to suggest DME might homodimerize to become active, any weak physical interaction caused by ectopic NTD expression might induce conformational change that renders DME non-functional. Unfortunately our attempt to assess whether the NTD of DME can interact with each other was confounded by the self-activating activity of DME.2 NTD in yeast two-hybrid assay when fused to the GAL4 DNA binding domain (data not shown). Their possible interaction will need to be assessed by alternative strategies. Another possibility is that NTD binds and titrates out an important interacting partner required to activate DME through conformational change (allosteric interaction). By removing NTD, the AGB region is liberated from such conformational constrain and can demethylate its target sites. It is also possible that the non-specific DNA binding activity of NTD competes with DME for target sites, thereby reducing the overall efficiency of DME. The molecular underpinning of how NTD induces DN effect remains to be elucidated. From an evolutionary viewpoint, the use of an active DME-based demethylation appears to have been acquired early in the plant lineage. The presence of several accessory domains in addition to the conserved core suggests adjustments to the chromatin and methylation environment of the different species. The presence of additional domains such as the DemeN and basic repeats in angiosperms and the permuted CXXC domain in streptophyta lineage might reflect the adjustment to the unique methylation and chromatin environment of the larger Streptophyta and land plant genomes.

## Materials and Methods

### Molecular Cloning of Constructs Used in this Study

All general molecular manipulations followed standard procedures (Sambrook et al. 1989). Q5 High Fidelity DNA polymerase (NEB, Ipswich MA, USA) was used for PCR amplifications. PCR products were purified using AMPure XP beads (Beckman Coulter, Indianapolis IN, USA). The sequences of all plasmid constructs were confirmed by sequencing (Eton, Research Triangle Park NC, USA). All PCR primers and double-stranded DNA fragments were synthesized by Integrated DNA Technologies (Coralville IA, USA), and sequences are listed in Supplementary File 1.

A binary plasmid vector, pFGAMh, was modified to facilitate the generation of plasmid constructs using the Gibson assembly method. In brief, the replication origins and T-DNA borders originated from pFGC5941 (GenBank Accession: AY310901). A hygromycin resistance gene (HPTII) under the control of the mannopine synthase promoter was installed for selection of transgenic seedlings. A Gateway attR cassette (rfa, Invitrogen, Carlsbad CA, USA), flanked with unique restriction sites XhoI and XbaI-SpeI was placed upstream octopine synthase polyadenylation signal (OCS3’). Plasmid pFGAMh, digested with restriction enzymes XhoI and XbaI, was used to generate plasmids pDME:DME^CTD^, pDME:nDME^CTD^ and pDME:GFP::DME^CTD^ using the Gibson assembly method. The DME.2 upstream regulatory sequence (DMEpro; 2895 bp upstream of DME.2 translation start codon ATG) was PCR-amplified using primer pair VeDME/P3R and Col-0 gDNA as template. The coding sequence of linker-AGB (with a 6-Ala linker to its N-terminus; 3174 bp), was PCR-amplified using primer pair lnAGBF/CTDVeR and Col-0 cDNA as template. To bridge these two fragments (DMEpro and linker-AGB), one of the following three DNA fragments was used in the assembly reactions. For pDME:DME^CTD^, a 50-bp fragment was generated by annealing DNA oligos ATGF and ATGR. For pDME:SV40NLS::AGB, a 71-bp fragment was generated by annealing DNA oligos S40F and S40R followed by two rounds of PCR reactions. For pDME:GFP::DME^CTD^, a 761-bp fragment was PCR-amplified using primer pair p3GFPF/dmGFPR and plasmid DNA pGFP-JS (Jen Sheen,Massachusetts GeneralHospital, Boston MA, USA) as template.

An intermediate plasmid vector, DME-P3-attR-AGB, was generated by digesting plasmid pDME:SV40NLS::AGB with restriction enzymes AflII and NcoI, and re-assembled with a 2800-bp fragment, which was produced through overlap PCR with 3 primer pairs, upAflII/P3attR, P3attF/attAGBR and attAGBF/dnNcoI, and Col-0 gDNA, attR cassette and Col-0 cDNA as templates. The resulting plasmid DME-P3-attR-AGB bears (1) the same 2895-bp regulatory sequence as the above constructs, (2) an attR cassette flanked by unique restrict sites XbaI and BglII, and (3) AGB coding sequence (3156 bp). To generate pDME:DME^FL^, plasmid DME-P3-attR-AGB was digested with XbaI and BglII, and assembled with a 2985-bp sequence, which was generated through overlap PCR using primer pairs S1-5e/IN3R and IN3F/S1-5R, and Col-0 gDNA as template. The resulting plasmid pDME:DME^FL^ carries the complete DME.2 coding sequence and intron 2 sequence (6075 bp) immediately downstream of the 2895-bp regulatory sequence with no additional sequences.

The intermediate plasmid vector DME-P3-attR-AGB was digested with restriction enzymes BglII and SpeI (to completely remove the AGB coding sequence), and re-assembled with a 786-bp sequence, which included the coding sequence of GFP (with its start codon ATG changed to TTG) and was PCR-amplified using primers ttGFPF and SpeGFPR and plasmid DNA pGFP-JS as template. The resulting plasmid DME-P3-attR-GFP was used as an intermediate plasmid vector to generate constructs pDME:DME^NTD^::GFP and pDME:mDME^NTD^::GFP. Plasmid DME-P3-attR-GFP was digested with XbaI and BglII, and assembled with two DNA fragments: a 3289-bp sequence was PCR-amplified using primers S1-5F and dme2tR2 and Col-0 gDNA as template and a 158-bp synthetic DNA fragment (FragQ20) (Integrated DNA Technologies, Coralville IA, USA). The resulting construct pDME:DME^NTD^::GFP included the 2895-bp upstream regulatory sequence, the 3332-bp sequence downstream of translation start codon ATG, the coding sequence of 6-Ala linker, and the coding sequence of GFP. Note the NTD coding sequence included the first 86 bp of intron 4 of gene DME.2, and it was designed to mimic dme-2 T-DNA insertion. To generate pDME:mDME^NTD^::GFP, the sequence of the first 1012 amnio acid residues of DME.2 protein was converted to DNA sequence using program EMBOSS Backtranseq (http://www.ebi.ac.uk/Tools/st/emboss_backtranseq/) and the Homo sapiens codon usage table. The sequence was then analyzed using online programs SoftBerry FSPLICE (http://linux1.softberry.com/berry.phtml?topic=fsplice&group=programs&subgroup=gfind) and NetPlantGene2 (http://www.cbs.dtu.dk/services/NetPGene/), and manually edited to disrupt potential splicing donor sites or acceptor sites. The mDME^NTD^ sequence (3036 bp) and upstream-and downstream-overlapping sequence are broken into 4 fragments, and synthesized by Integrated DNA Technologies (Coralville IA, USA). The 4 DNA fragments were assembled with plasmid DME-P3-attR-GFP digested with XbaI and BglII, resulting construct pDME:mNTDh::GFP.

### Whole-Genome Bisulfite Sequencing and DNA Methylome Analysis

Genomic DNA were isolated from hand dissected, 7-9 DAP *dme-2* endosperm that has been complemented by *DME*^*FL*^ or *nDME*^*CTD*^ (*dme-2*/*dme-2*;*DME*^*FL*^*/DME*^*FL*^ or *dme-2/dme-2; nDME*^*CTD*^ *nDME*^*CTD*^). Whole genome bisulfite sequencing library was constructed as described before ^13, 44^. Approximately 20-50 ng of purified genomic DNA was spiked with 0.5ng of unmethylated cl857 *Sam7* Lambda DNA (Promega, Madison, WI) and sheared to about 300bp using Covaris M220 (Covaris Inc., Woburn, Massachusetts) under the following settings: target BP, 300; peak incident power, 75 W; duty factor, 10%; cycles per burst, 200; treatment time, 90 second; sample volume 50μl. The sheared DNA was cleaned up and recovered by 1.2x AMPure XP beads then followed by end repaired and A-tailing (NEBNext Ultra II DNA Library Prep Kit for Illumina, NEB) before ligation to NEBNext methylated multiplex adapters (NEBNext Multiplex Oligos for Illumina, NEB) according to the manufacturer’s instructions. Adaptor-ligated DNA was cleaned up with 1x AMPure XP beads. The purified adaptor-ligated DNA was spiked with 50ng of unmethylated cl857 *Sam7* Lambda DNA and subjected to one round of sodium bisulfite conversion using the EZ DNA Methylation-Lightning Kit (Zymo Research Corporation, Irvine, CA) as outlined in the manufacturer’s instructions with 80 min of incubation time. Half of the bisulfite-converted DNA molecules was PCR amplified with the following condition: 2.5 U of ExTaq DNA polymerase (Takara), 5 ul of 10 x Extaq reaction buffer, 25 μM dNTPs, 1 ul of index Primers (10 uM) in 50 uL reaction. The thermocyling condition was as follows: 95 °C for 2 min and then 10 cycles each of 95 °C for 30 s, 65 °C for 30 s, and 72 °C for 60 s. The enriched libraries were purified twice with 0.8x (v/v) AMPure XP beads to remove any adapter dimers. High throughput sequencing was performed by Novogene Corporation (USA). For each genotype, sequencing reads from three individual transgenic lines were combined. Sequenced reads were mapped to the TAIR10 reference genomes and DNA methylation analyses were performed as previously described (Supplementary Table 7) ^13^. Fractional CG methylation in 50-bp windows across the genome was compared between *dme*, wild-type (GSE38935 ^13^), *DME*^*FL*^*-nDME*^*CTD*^*-* complemented *dme-2* endosperm. Windows with a fractional CG methylation difference of at least 0.3 in the endosperm comparison (Fisher’s exact test p-value < 0.001) were merged to generate larger differentially methylated regions (DMRs) if they occurred within 300 bp. DMRs were retained for further analysis if the fractional CG methylation across the whole DMR was 0.3 greater in *dme* endosperm than in wild-type endosperm (Fisher’s exact test p-value < 10^-10^), and if the DMR is at least 100-bp long. The merged DMR lists are in the Supplemental File 2. The *dme* and wild-type endosperm data used in this study were derived from crossed between *Col* (female parent) and *Ler* (male parent) (GSE38935, ^13^). To avoid potential ecotype-specific methylation difference, *Ler* hyper-DMRs relative to Col-0 endosperms (GSE52814, ^45^) were identified using the same criteria as described above and excluded from further analyses. For making the Venn diagram, merged DMR regions were converted into 50-bp windows. Only windows with methylation scores in all samples were retained for comparison in Venn diagram and boxplot analysis.

### Plant Materials and Complementation Assays

We found we can easily obtained *dme-2/dme-2* Col-*gl* plants from *DME/dme-2* heterozygotes if we rescued seeds prior to desiccation on MS sucrose plates. This is consistent with the report that *fis* endosperm cellularization defect and embryo arrest can be rescued by culturing the developing seeds in sucrose media because *fis* seeds have reduced hexose level ^46^. Using this method we generated multiple homozygous lines, and we did not detect any difference between individuals in terms of normal seed rate or visible phenotype. The adult *dme-2/dme-2* plants are morphologically indistinguishable from wild-type Col-*gl* plants but produce ∼0.1% viable mature seeds. These *dme-2/dme-2* plants are not due to genetic mutation or heritable aberrant epigenetic effects that escape requirement of DME activity during gametogenesis because their subsequent progeny are phenotypically normal and produces same level (∼0.1%) of normal seeds.

The *DME/dme-2* heterozygous or *dme-2/dme-2* homozygous lines in Col-*gl* background were subjected to Agrobacterium-mediated floral dipping transformation procedures ^28^. Seeds were sterilized by 30% bleach solution and screened for T1 transgenic plants on a 0.5x MS nutrient medium with 1% sucrose, 0.8% agar and 40 μg/ml hygromycin. Germinated seedlings were transferred to soil and grown in the growth room under 16 hours of light and 8 hours of dark cycles at 23°C. Siliques from T1 transgenic plants were dissected 14-16 days after self-pollination using a stereoscopic microscope (SteREO Discovery.V12, Carl Zeiss, Wetzlar, Germany). The numbers of viable and aborted seeds in transgenic lines were statistically analyzed with the χ^2^ test. The probability that deviates from a 1:1 or 3:1 segregation ratio for viable and aborted seeds was also calculated.

### RNA extraction, cDNA synthesis and quantitative PCR analysis

Total RNA was extracted using TRIzol(®) Reagent (Invitrogen, Carlsbad, USA) and treated with TURBO DNase (Ambion, Austin TX, USA) according to the manufacturers’ instructions. For cDNA synthesis, 5mg of total RNA was reverse-transcribed using Superscript III Reverse Transcriptase and oligo(dT) primer (Invitrogen). The cDNA was treated with RNase H (Invitrogen) at 37oC for 20min and diluted tenfold with H2O. For each 15-μl qPCR reaction, 1μl of diluted cDNA was used. The quantitative PCR was run on ABI 7500 Fast Real-Time PCR System (http://www.appliedbiosystems.com) using FastStart Universal SYBR Green Master Mix (Roche, http://www.roche.com). The quantitative PCR primers are listed in Supplementary File 1. The Ct values were normalized against *ACT2* (*At3g18780*) mRNA or *UBC* (*At5g25760*) mRNA. The abundance of mRNAs was expressed as relative to controls, with control values set to 1. The error bars represent the standard deviation of 4 biological replicates.

### Protein domain analysis and phylogenetic inference

We utilized a domain-centric computational strategy to study DME and its related proteins. Specifically, we identify DME homologs by using the iterative profile searches with PSI-BLAST47 from the protein non-redundant (NR) database at National Center for Biotechnology Information (NCBI). Multiple sequence alignments were built by the Promals ^48^ program, followed by careful manual adjustments. Consensus secondary structures were predicted using the PSIPRED ^49^ JPred program ^50^. Conserved domains were further characterized based on the comparison to available domain models from pfam ^51^ and sequence/structural features. The PhyML program ^52^ was used to determine the maximum-likelihood tree using the Jones–Taylor– Thornton (JTT) model for amino acids substitution with a discrete gamma model (four categories with gamma shape parameter: 1.096). The tree was rendered using MEGA Tree Explorer ^53^.

## Acknowledgments

We thank Drs. Ping-Hung Hsieh and Jer-Young Lin for assistance in methylome analysis, Robert Goldberg, Robert Fischer and Matthew Bauer for discussions during the course of this study. This work is partially supported by the Hatch Project 02413 (to T.-F.H.), National Institute of Food and Agriculture, U.S. Department of Agriculture, U.S.A., by the National Science Foundation Grant MCB-1715115 to T.-F.H. and W.Y.X., and by the State of NC appropriations as distributed by the University of North Carolina General Administration and the NC Agricultural Research Service Office at NC State University. LMI and LA are supported by the Intramural Research Program of the National Library of Medicine, NIH, USA.

## Author contribution.

C.Z., Y.-H.H., X.-Q.Z., J.H.H. and T.-F.H. designed the research. C.Z., Y.-H.H., X.-Q.Z. performed the experiments. D.Z., L.M.I, and L.A. performed the evolutionary analysis. C.Z., Y.-H.H., and T.-F.H. wrote the article. T.-F.H., C.Z., Y.-H.H., W.X., J.H.H. interpreted and commented the article.

## Supplemental Information

**Fig. S1.**
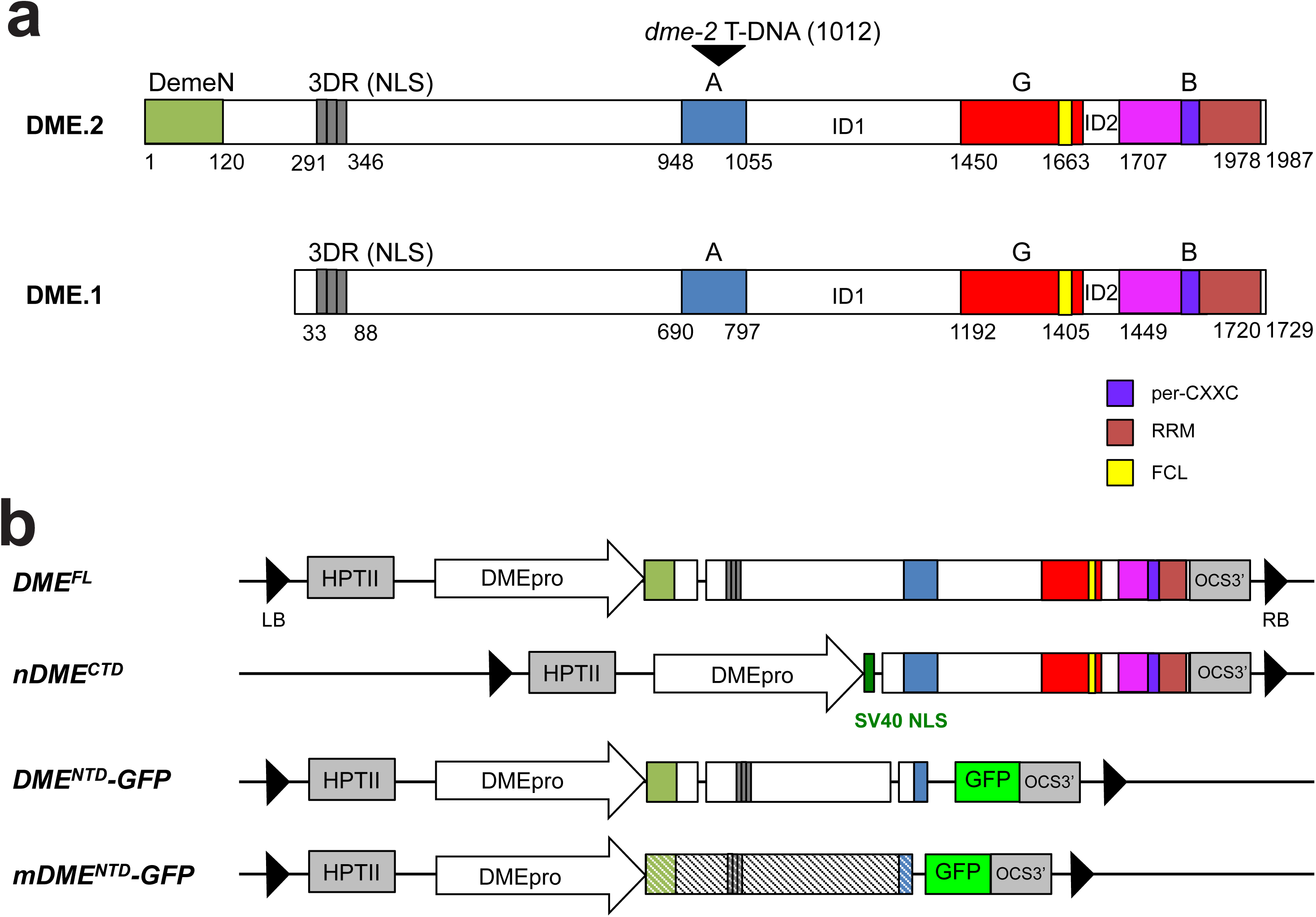
Diagrams of DME protein structure and transgene constructs. (a) DME protein domain architecture. The positions of conserved domains along DME protein. Numbers represent amino acid position relative to the translation start sites. DME.1 is shorter than DME.2 by 258 amino acids due to alternative splicing, missing the very N-terminal DemeN domain. DemeN is a domain of unknown function conserved among angiosperm DME-like protein. 3DR is the stretch of basic rich amino acid direct repeats, resembling AT-hook motifs, and serves as a nuclear localization signal; per-CXXC is the permuted CXXC zinc finger motif; RRM is the RNA recognition motif; FCL is a [Fe4S4] cluster loop following the HhH module. The *dme-2* allele harbors a T-DNA insertion in region A at amino acid position 1012. ID1 and ID2 are variable, low complexity sequences between the glycosylase domain and the conserved B region. (b) Transgene constructs used in this study. DMEpro refers to the upstream regulatory sequence (2895 bp upstream of the translation start codon ATG) of DME.2. SV40NLS: PKKKRKV. A polypeptide linker comprising 6 alanine residues is placed between any protein fragment fusions.

**Fig. S2.**
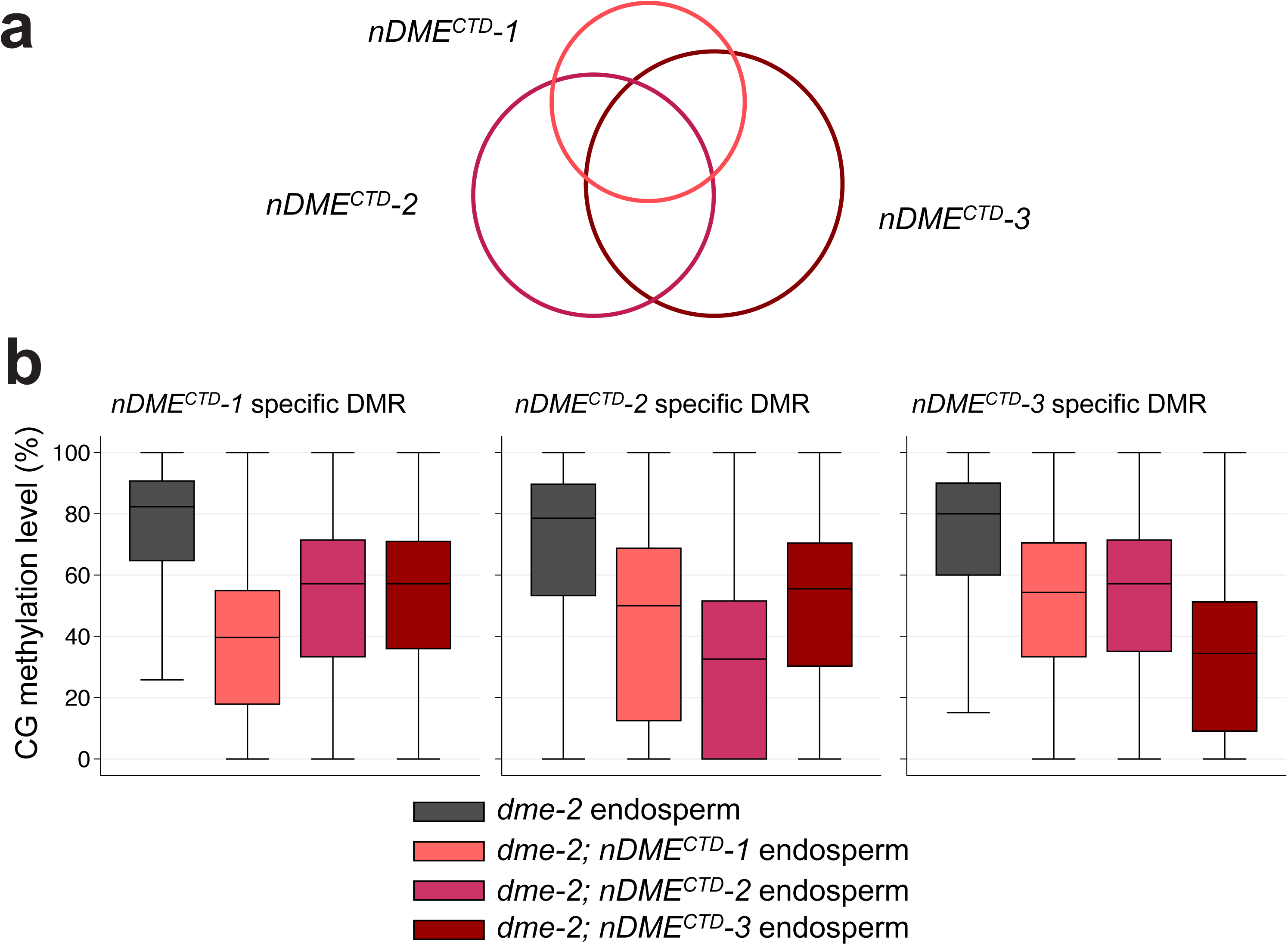
DNA methylomes of three independent *nDME*^*CTD*^-complemented*dme-2* endosperm. (a) Venn diagram showing partial overlap of *dme* CG hyper-DMRs relatives to each nAGB-complemented endosperm (*nDME*^*CTD*^*-1* to *nDME*^*CTD*^-3). (b) Boxplot of CG methylation levels among canonical DME target sites in *dme-2* mutant (black), *nDME*^*CTD*^-1 (pink), *nDME*^*CTD*^*-2* (magenta), or *nDME*^*CTD*^*-3* (red) complemented endosperm, in *nDME*^*CTD*^*-1* specific (left panel), *nDME*^*CTD*^*-2* specific (middle panel), and *nDME*^*CTD*^*-3* specific DMRs. These results show that the combined DMRs are more or less hypomethylated in each independent line compared to *dme-2* endosperm.

**Fig. S3.**
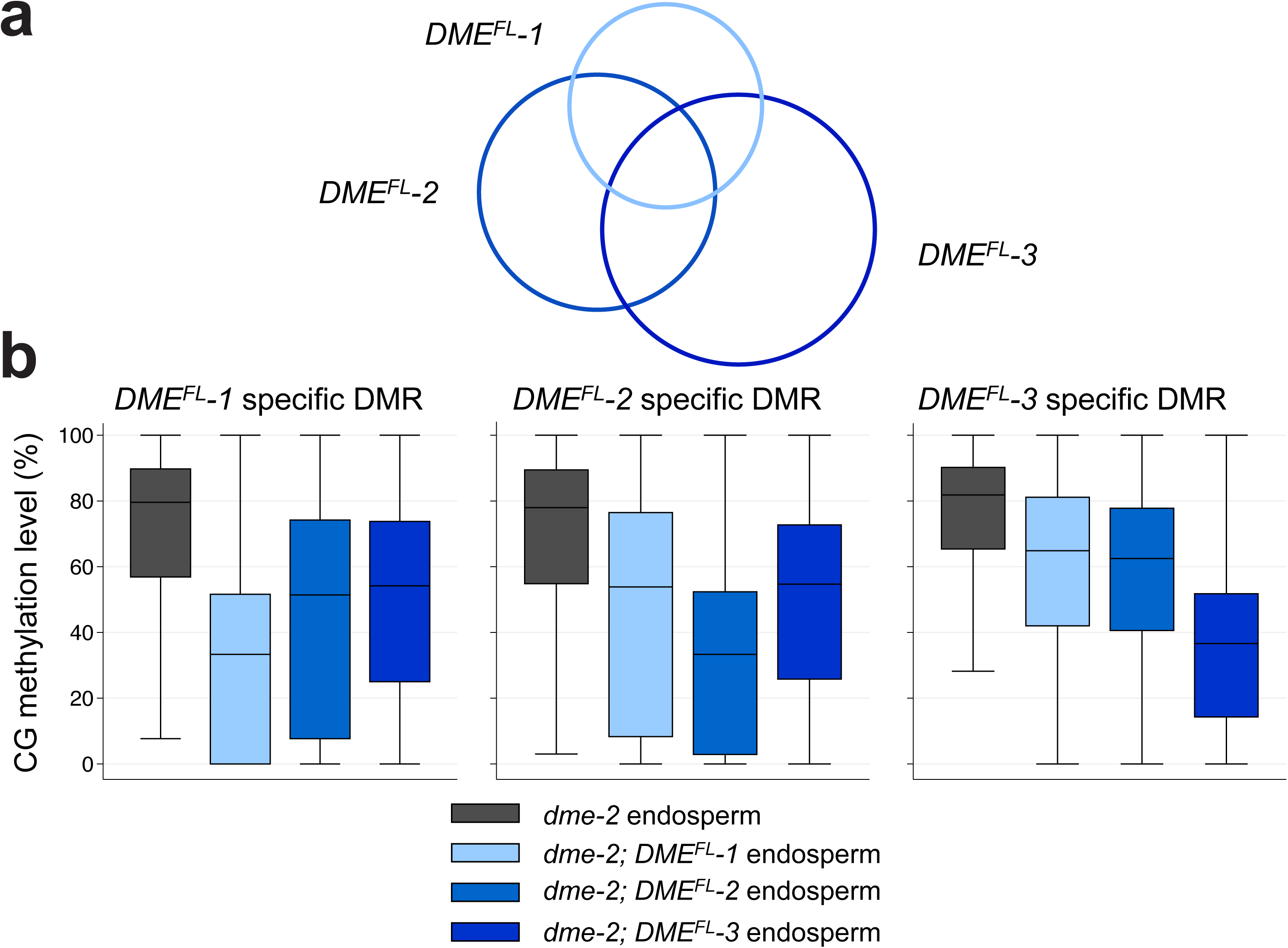
DNA methylomes of three independent DME^FL-^complemented *dme-2* endosperm. (a) Venn diagram showing partial-overlap of *dme* CG hyper-DMRs relatives to each *DME*^*FL*^-complemented endosperm (*DME*^*FL*^-*1* to *DME*^*FL*^-*3*). (b) Boxplot of CG methylation levels among canonical DME target sites in *dme-2* mutant (black), *DME*^*FL*^-1 (light blue), *DME*^*FL*^-*2* (medium blue), or *DME*^*FL*^-*3* (dark blue) complemented endosperm, in *DME*^*FL*^-*1* specific (left panel), *DME*^*FL*^-*2* specific (middle panel), and *DME*^*FL*^-*3* specific DMRs. These results show that the combined DMRs are more or less hypomethylated in each independent line compared to *dme-2* endosperm.

**Fig. S4.**
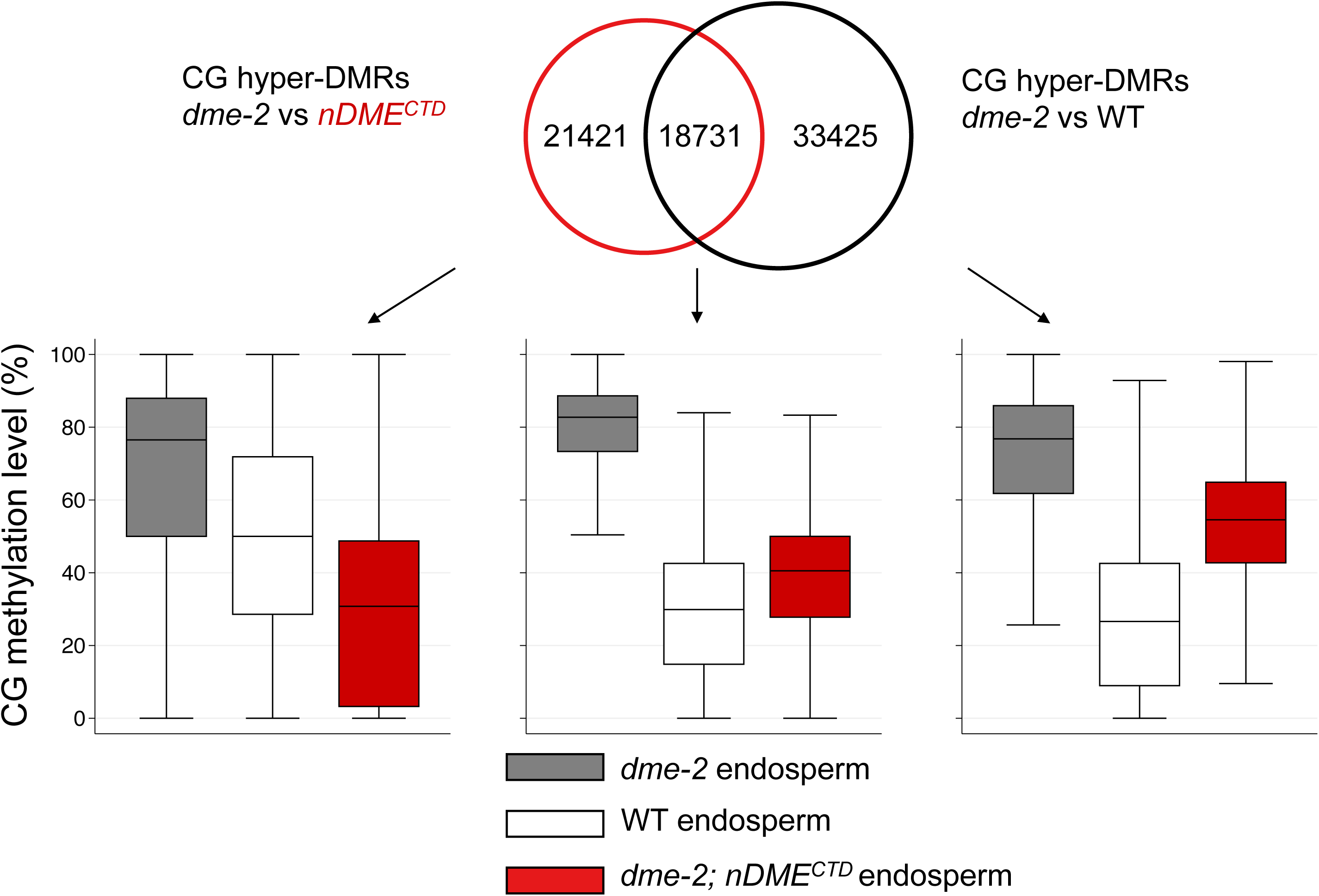
The DMRs of *dme* relative to WT endosperm or nDME^CTD^-complemented endosperm. Venn Diagram (top) and Boxplot analysis (bottom) of CG hyper-DMRs in 50-bp windows between *dme-2* endosperm relative to *nDME*^*CTD*^-complemented or WT endosperm. CG methylation levels of DMRs unique to *nDME*^*CTD*^-complemented endosperm are also demethylated in the WT endosperm (left panel). Similarly, DMRs unique to WT endosperm are demethylated in *nDME*^*CTD*^-complemented endosperm (right).

**Fig. S5.**
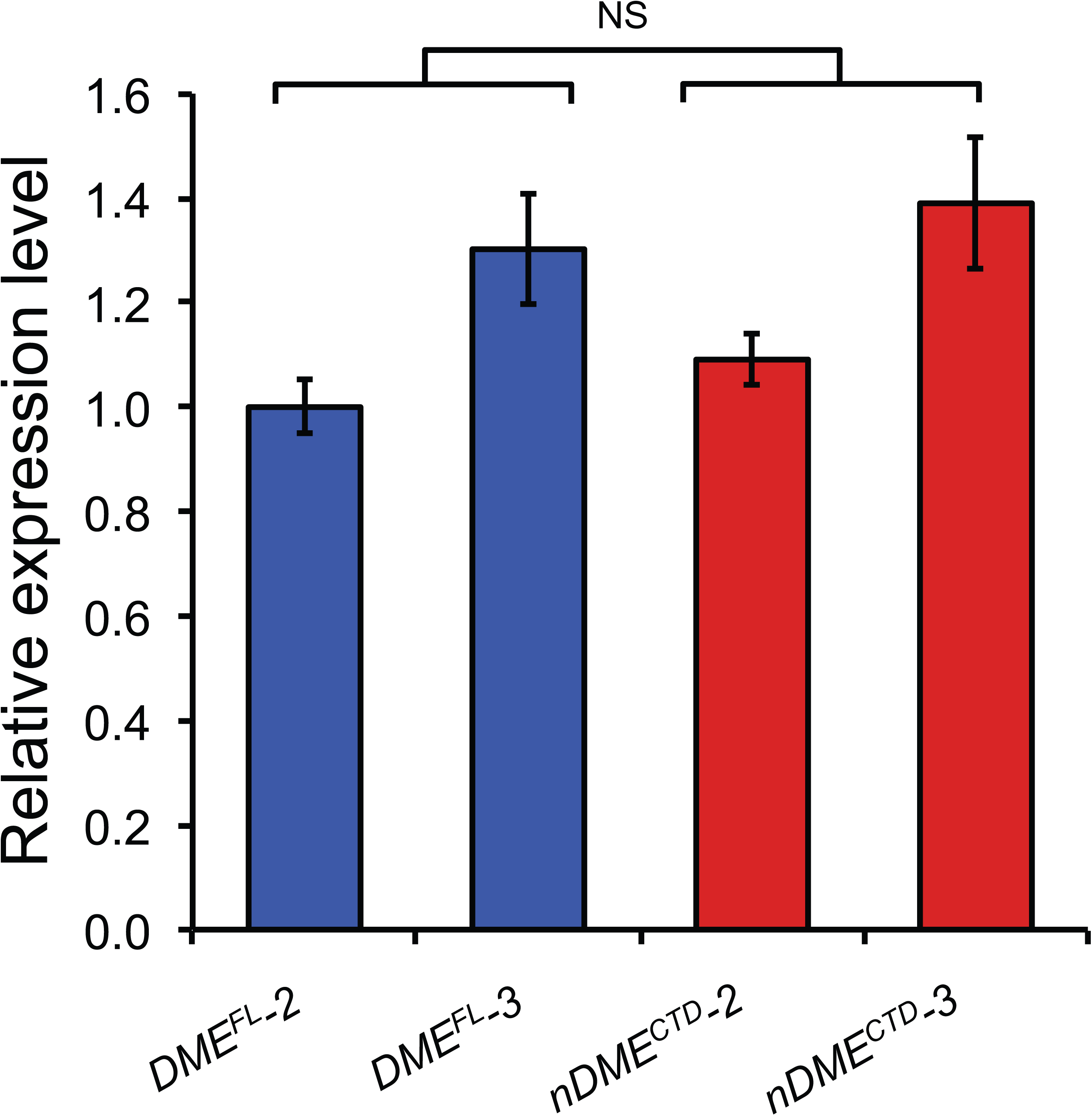
DME^FL^ and nDME^CTD^ transgenes are expressed at comparable levels among independent complementation lines. *DME*^*FL*^ and *nDME*^*CTD*^ expression levels are comparable between the four of the six complementation lines used in the methylome study. Total RNA was isolated from stage F1 to F12 floral buds. The results show that there is no significant difference in expression level between these two transgenes(t-test, *p*>0.4).

**Fig. S6.**
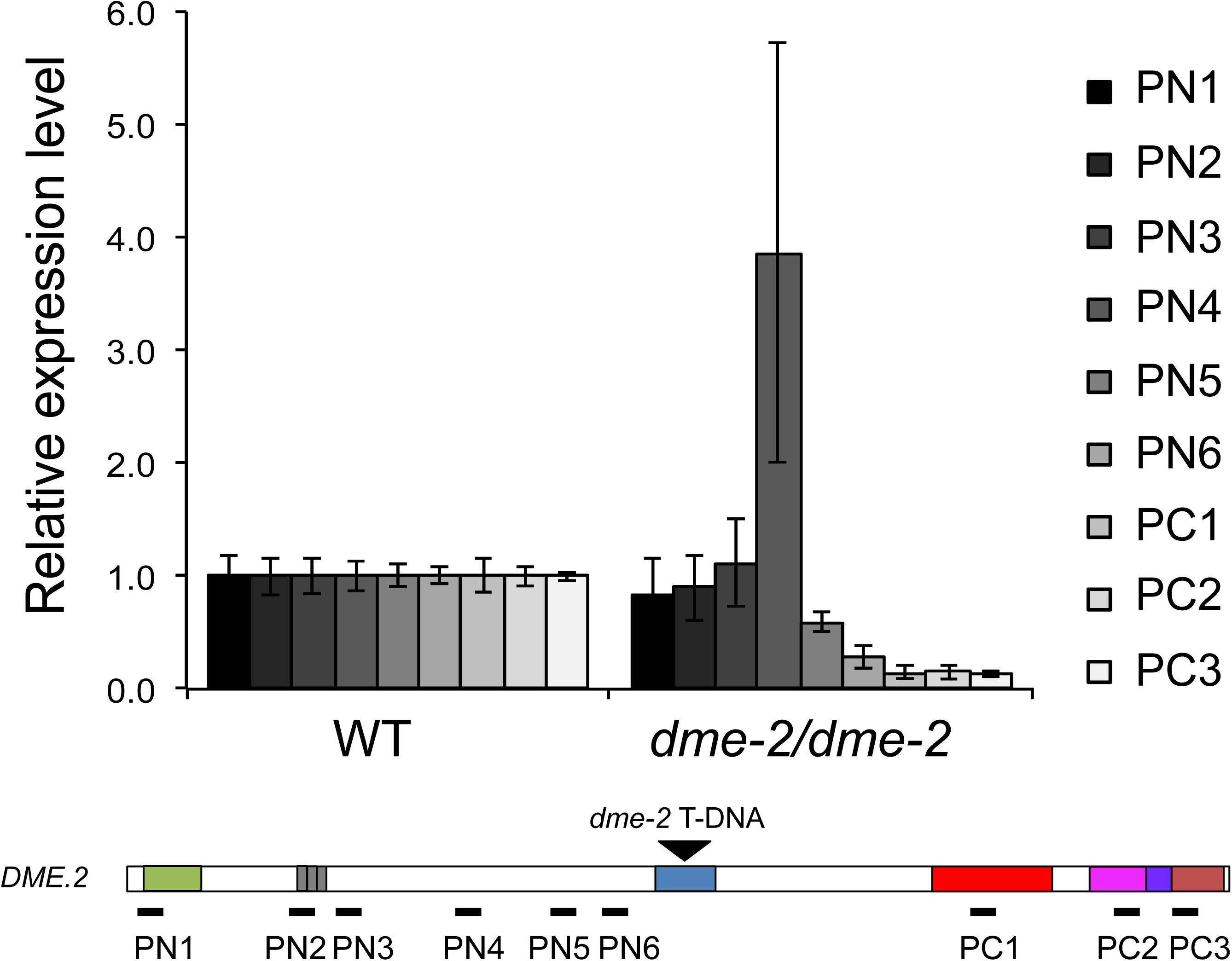
The effects of T-DNA insertion on endogenous DME transcript abundance in *dme-2/dme-2* plants. Total RNA was isolated from stage F1 to F12 floral buds. Equal amount of total RNA from WT and *dme-2/dme-2* were used for reverse transcription and quantitative PCR. Six paired of primers (PN1-PN6) correspond to the N-terminal region before the T-DNA insertion site, and three pairs of C-terminal region primers (PC1-PC3) were used to assess endogenous DME transcript level in *dme-2/dme-2* mutant plants. The position of each primer pair is indicated in the DME diagram where T-DNA insertion site is shown.

**Fig. S7.**
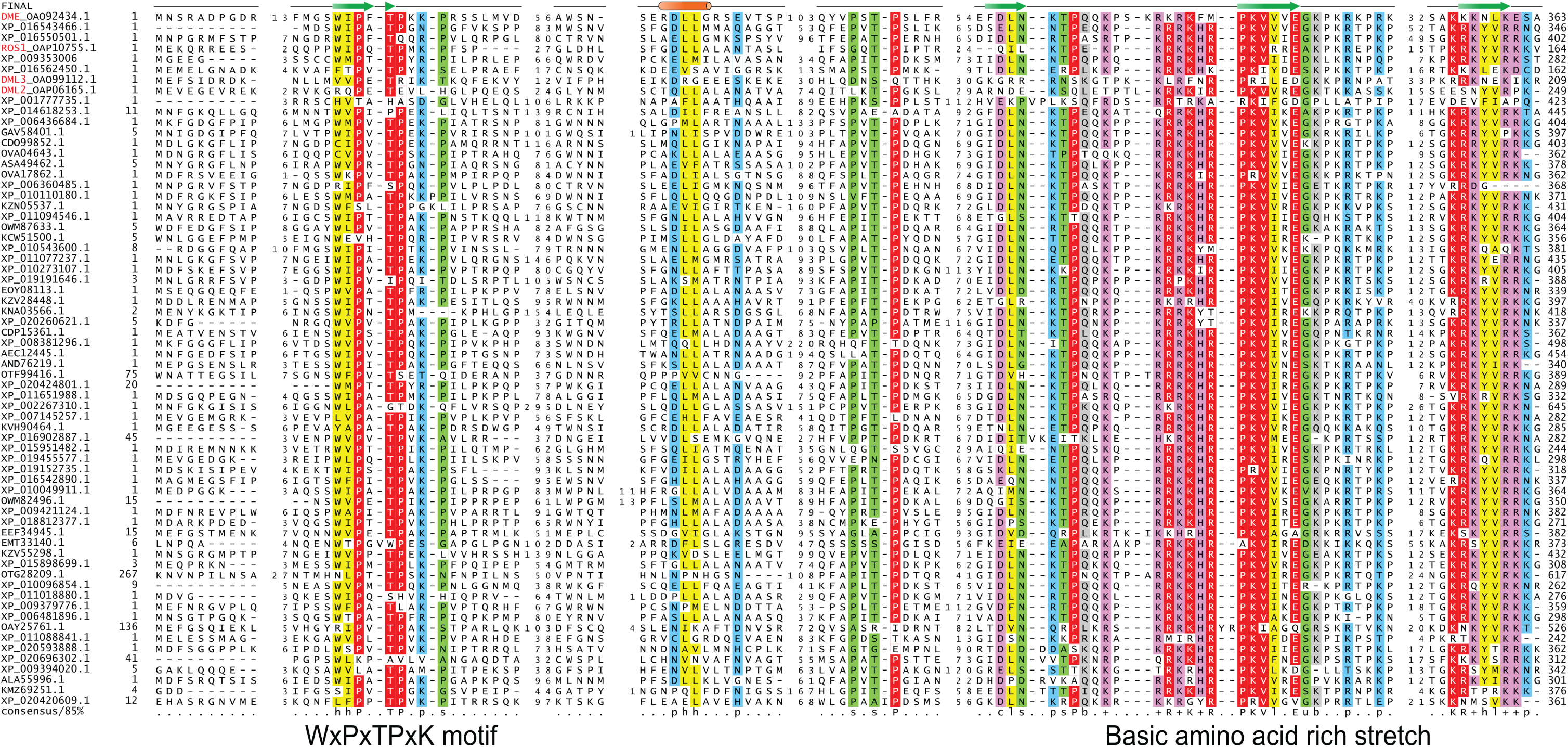
Alignment of angiosperm DME-like proteins showing the conserved DemeN domain and the basic rich 3DR repeats. Bioinformatics analysis using available DME-like sequences identified a ∼ 120-amino-acid-long conserved region at the very N-termini among DME-like proteins in angiosperms. This sequence is characterized by a highly conserved WxPxTPxK motif that might function in protein-protein interactions. Further toward the C-terminus is a stretch of basic amino acids rich region that serves as a nuclear localization signal. This sequence consists of three direct repeats (3DR) reminiscent of the AT-hook motifs that may bind DNA.

**Fig. S8.**
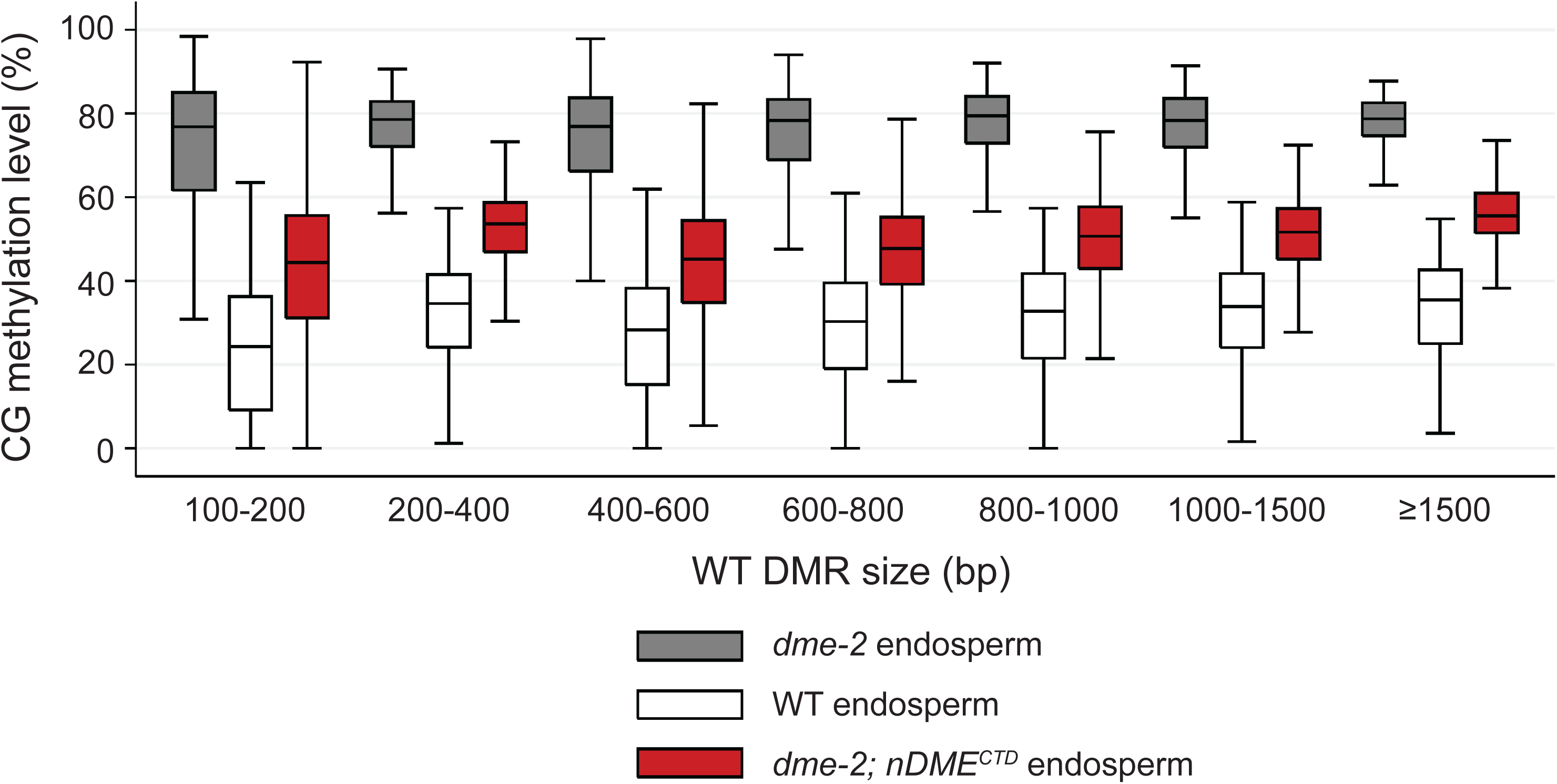
Boxplot of CG methylation levels among canonical DME target sites in different DMR length category, in *dme-2* mutant (black), wild-type (white), or *nDME*^*CTD*^-complemented endosperm

**Fig. S9.**
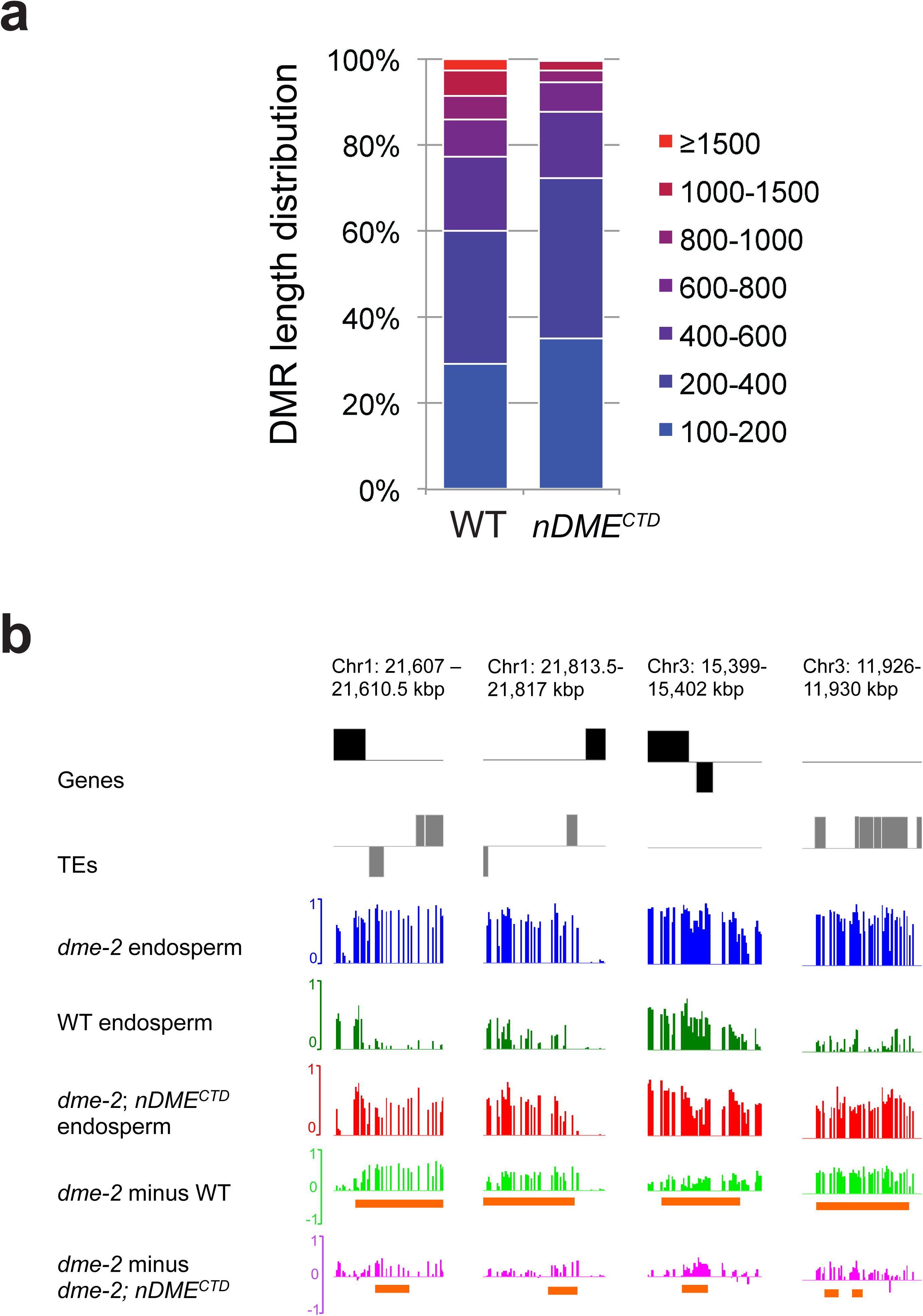
(a) Merged DMR length distribution in WT and *nDME*^*CTD*^-complemented endosperm. (b) Genome Browser examples of long WT DMRs. Tracks are as labeled. The DMR regions are indicated as horizontal bars according to their length in each sample (bottom two tracks). Even though *nDME*^*CTD*^ complemented endosperm lack longer DMRs, these regions are also shorter DMRs in *nDME*^*CTD*^-complemented endosperm.

**Table 1.**
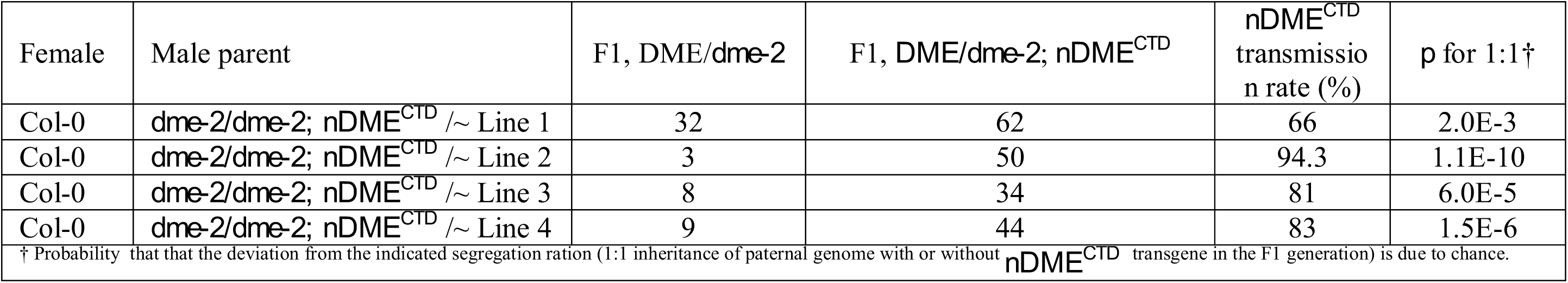
Rescue of the reduced paternal *dme-2* allele transmission by the *nDME*^*CTD*^ transgene.

